# Increased Low Molecular Weight Mucins in Muco-Obstructive Airway Disease Limit *Staphylococcus aureus* Growth

**DOI:** 10.64898/2025.12.03.692258

**Authors:** Caitlyn C. Sebastian, RaNashia Boone, Susan E. Birket, Megan R. Kiedrowski

**Author notes:** Address correspondence to Megan R. Kiedrowski.

## Abstract

Muco-obstructive airway diseases result in an increase in mucus accumulation and a decrease in mucus clearance. MUC5B is the most abundant secreted mucin in the human airways, and MUC5B mucin strands dimerize to create the mucus mesh network in the healthy respiratory tract. In muco-obstructive airway diseases like cystic fibrosis (CF), immune cells and bacteria release enzymes that degrade MUC5B into smaller fragments that become entangled and compacted, contributing to pathogenesis. We utilized synthetic cystic fibrosis sputum media (SCFM) to examine how mucin polymers can impact *Staphylococcus aureus*, a common CF pathogen that persists despite highly effective modulator therapies to correct CF disease. We found low molecular weight (LMW) mucin negatively impacts *S. aureus* survival and biofilm biomass compared to high molecular weight (HMW) mucin. Adding extracellular DNA to SCFM with LMW mucin was not sufficient to restore growth. LMW mucin had a broad negative impact on *S. aureus* laboratory strains and CF clinical isolates. We next tested other CF pathogens, including *Pseudomonas aeruginosa* and nontypeable *Haemophilus influenzae*, and saw no significant differences in growth in HMW or LMW mucin. LMW mucin did not significantly impact *Staphylococcus epidermidis* growth, indicating there may be specific interactions with *S. aureus*. Overall, this work highlights how interactions with pathogenic mucins may limit *S. aureus* growth in the diseased airways while supporting low-level persistence, and its ability to thrive in the presence of longer mucin strands may help to explain why *S. aureus* is well adapted to survive in the healthy respiratory tract.

**IMPORTANCE:** *Staphylococcus aureus* is a common colonizer of the healthy human airways that is also known to establish persistent infections in the lungs of people with muco-obstructive airway diseases, including cystic fibrosis. Yet, few studies have investigated how *S. aureus* interactions with airway mucins in chronic airway disease may regulate bacterial persistence or pathogenesis. In this study, we report that low molecular weight mucins representative of short, degraded mucin polymers found at high abundance in muco-obstructive airway disease suppress *S. aureus* growth and limit bacterial biofilm formation and aggregation. Longer mucin polymers did not negatively affect *S. aureus* survival. Other common CF pathogens were not significantly affected by low molecular weight mucins, suggesting there are species-specific interactions between airway mucins and *S. aureus* in the chronically diseased lung. This work highlights how pathogenic mucins found in muco-obstructive disease can affect *S. aureus* growth and promote phenotypes associated with airway persistence.

## INTRODUCTION

Muco-obstructive airway diseases, such as cystic fibrosis (CF), primary ciliary dyskinesia (PCD), and chronic obstructive pulmonary disease (COPD), are characterized by decreased mucociliary clearance and increased mucus accumulation in the airways (1, 2). The respiratory tract mucosal barrier contains heavily glycosylated tethered and secreted mucins that prevent adhesion of foreign particles and provide scaffolding for antibodies and other antimicrobial materials (3). In muco-obstructive diseases, secreted mucins become compacted and entangled due to hypersecretion and dehydration, which leads to abnormalities in mucin folding (2, 4). Of the secreted mucins, MUC5B is the most abundant in human airways (5). MUC5B is secreted from submucosal glands as linear strands and elongated through dimerization of disulfide linkages, creating the mucus mesh network (6–8). However, in muco-obstructive diseases including CF, mucin degradation is commonly observed due in part to persistent inflammation and infection. Bacteria and immune cells in the diseased airways contribute to mucin degradation through the release of proteases, resulting in an increased abundance of short MUC5B strands and compromising the function of the mucus layer (4, 9, 10). Compacted stagnant mucin creates a unique niche that allows for bacteria to access nutrients resulting from degraded airway proteins and use the compacted mucin mesh as scaffolding for aggregation and biofilm formation (1, 11–13).

*Staphylococcus aureus* is a common respiratory pathobiont that asymptomatically colonizes the upper respiratory tract of approximately 40% of healthy adults (14, 15). High prevalence of *S. aureus* has been reported in people with PCD and CF and is associated with higher risk of severe disease outcomes (16, 17). More recent studies indicate that increased prevalence of *S. aureus* leads to higher mortality in people with COPD (18). *S. aureus* is known to bind to mucin strands and can utilize products from degraded mucins broken down by other bacteria in the upper respiratory tract (19, 20). In CF, *S. aureus* remains prevalent in the airways even after treatment with highly effective modulator therapies (HEMT), with continued *S. aureus* colonization observed up to three years post-HEMT treatment (21, 22). The use of HEMT potentiates functional cystic fibrosis transmembrane conductance regulator (CFTR) ion channels, which restores airway surface liquid hydration and allows for clearance of mucus out of the airways (21, 23). Together with microbiological data, this suggests *S. aureus* is adapted to persist in both the diseased airways in the presence of high mucin concentrations and in airways with restored mucociliary clearance. However, how interactions with mucins impacts *S. aureus* biogeography and persistence in the mucus obstructed airways has not been explored in great depth.

Synthetic cystic fibrosis sputum media (SCFM) mimics the nutritional environment of CF and can include extracellular DNA (eDNA) and mucin polymers that can be added in different concentrations to replicate the increased eDNA and mucin concentrations found in muco-obstructive airway diseases (1, 12, 24, 25). Bacterial transcriptomes in SCFM have been shown to be comparable to ex vivo CF sputum collected from persons with CF (pwCF) and other commonly used CF models (26, 27). While there are currently no widely available synthetic defined media models for other muco-obstructive airway diseases, there is evidence that of similar changes in metabolic profiles in the respiratory tract, such as an increase in ammino acids, lactate, phospholipids and sialic acids, of people with muco-obstructive airway disease compared to healthy individuals (28, 29).

Our work aimed to investigate how *S. aureus* interactions with mucins affect persistence in the CF airway environment. In this study, we used SCFM with different mucin polymer sizes to determine how smaller mucin strands impact *S. aureus* compared to longer strands, mimicking degraded pathogenic mucins found in disease and healthy airway mucins. We found that shorter mucin strands had a negative impact on *S. aureus* survival and biomass compared to longer mucin polymers for laboratory strains and clinical isolates, and reducing the concentration of smaller mucins partially recovered *S. aureus* survival and biofilm. Additionally, we did not see a negative effect of small mucin polymers on other CF pathogens tested including *Pseudomonas aeruginosa* or nontypeable *Haemophilus influenzae* (NTHi). When we evaluated survival of another *Staphylococcus* species, *Staphylococcus epidermidis,* we did not see a decrease in survival in the presence of small mucin polymers, indicating that this phenotype may be specific to *S. aureus*.

## RESULTS

### Low molecular weight mucin impedes *S. aureus* growth and biomass

To determine the impact of long and short mucin strands, we utilized two different commercially available bovine submaxillary mucins (BSM) of different molecular masses to represent a low molecular weight (LMW; 200-500 kDa) and high molecular weight (HMW; 400-4,000 kDa) mucin environment in CF. Transmission electron microscopy confirmed longer mucin polymers were present in the HMW mucin samples, but lacking in the LMW samples (Figure 1A and B). We inoculated a well characterized community-associated strain of methicillin-resistant *S. aureus* (MRSA) USA300 into SCFM without polymers and SCFM containing LMW or HMW mucin. We found that there was no significant difference in *S. aureus* colony forming units (CFU) after 24 hours of incubation in SCFM alone or SCFM with HMW mucin, however there was significant decrease in USA300 survival in SCFM with LMW mucin (Figure 1C). Using GFP-expressing USA300, we next examined how LMW and HMW mucins impact USA300 biofilm formation. We found a significant decrease in overall biomass in the LMW mucin condition compared to the HMW mucin condition (Figure 1D and E). Together, these results indicated that LMW mucins have a detrimental effect on USA300.

**Figure 1:**
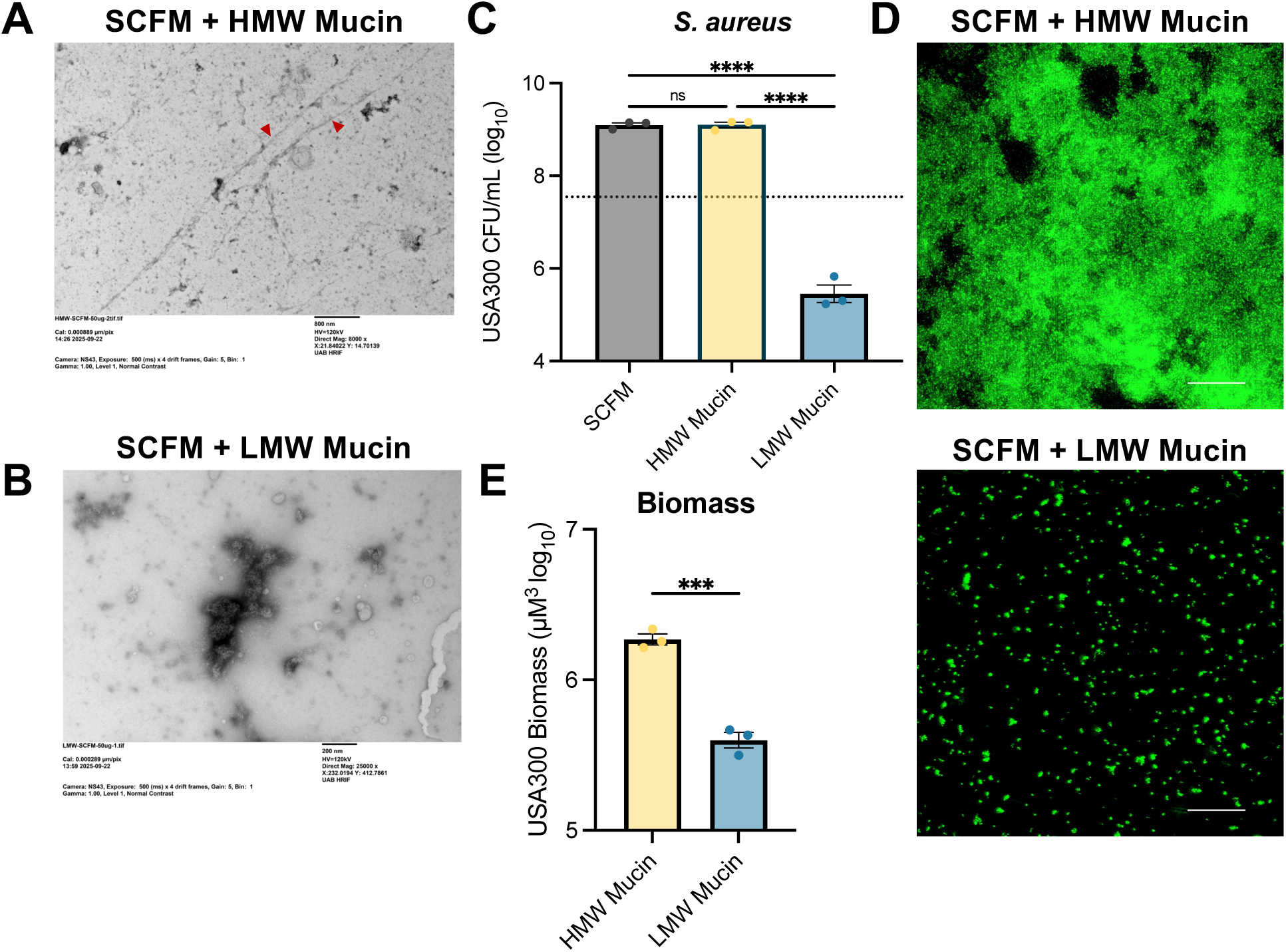
Low molecular weight mucins reduce *S. aureus* USA300 survival and biomass. Transition electron microscopy of uninfected (A) HMW mucin or (B) LMW mucin. Red arrows indicate long mucin polymer strands. USA300 growth (C) in SCFM (grey), HMW mucin (yellow) or LMW mucin (blue) for 24 hours, shaking. Dotted line represents the average inoculum into conditions. Representative images of GFP-expressing USA300 (D) in HMW mucin (top) or LMW mucin (bottom) z-stack images taken at 24 hours. Quantification of biomass (E) is represented by averaging 5 fields of views from 3 biological replicates for each condition. Scale bar represents 50 µm. One-way ANOVA was performed on USA300 colony-forming units (CFUs) and a Weltch’s t-test was performed on average biomass. (***P < 0.001, ****P < 0.0001). Data is represented by mean ± SEM.

To determine how LMW mucins impacted USA300 over time, we inoculated USA300 into SCFM containing LMW mucin and measured CFUs at 4-, 8-, 12- and 24 hours post-inoculation. We also inoculated USA300 into SCFM containing eDNA, both with and without LMW mucin, to determine if eDNA could rescue USA300. At 4 hours, we already saw a significant decrease in USA300 CFUs in SCFM with LMW mucin and SCFM with LMW mucin and eDNA compared to conditions where LMW mucins were not present (Figure 2B). We continued to see this significant decrease at later 8-, 12- and 24-hour time points. There remained no significant difference between SCFM with LMW mucin and SCFM with both LMW mucin and eDNA, indicating eDNA is not sufficient to rescue USA300 growth in the presence of LMW mucin.

**Figure 2:**
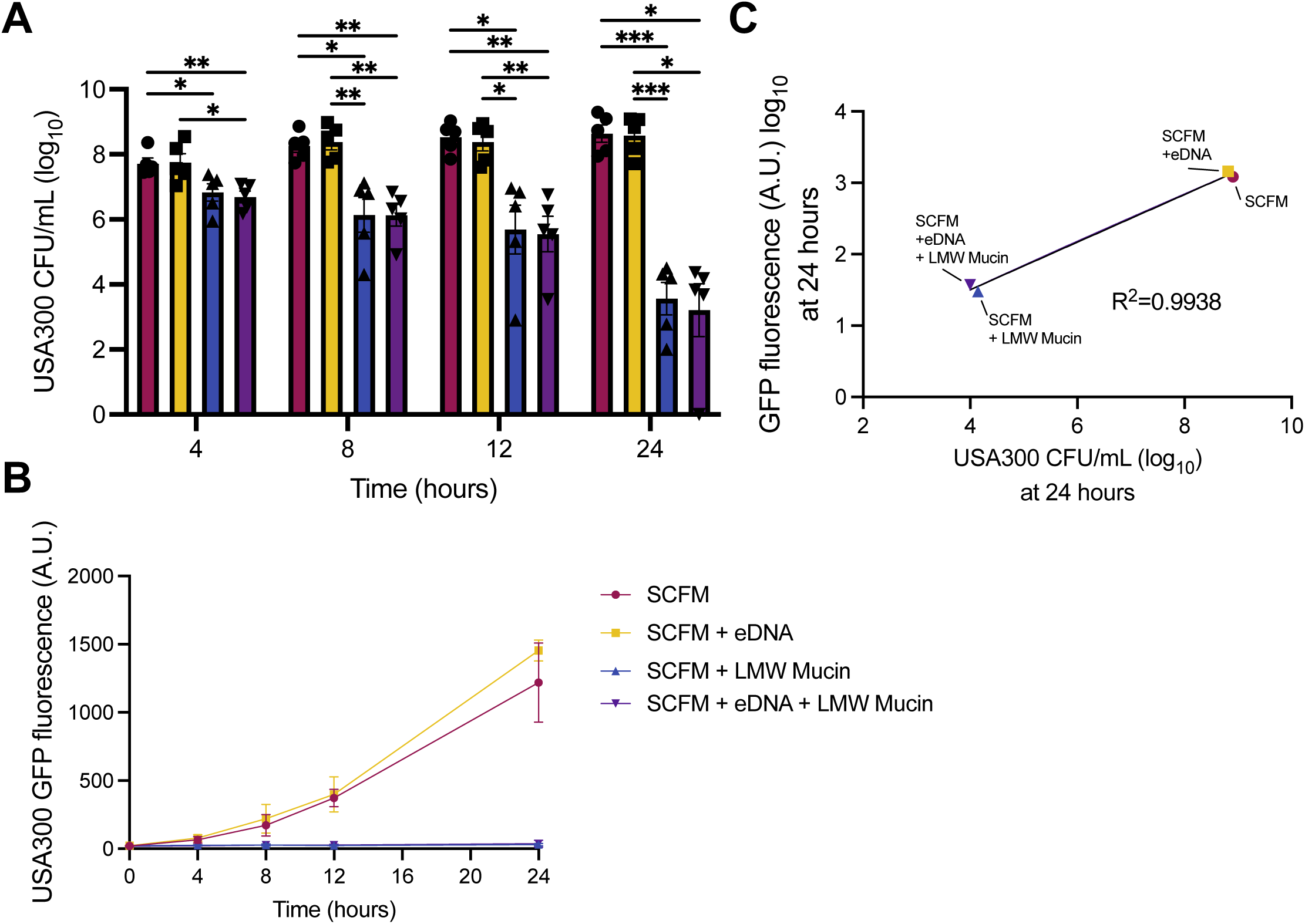
Low molecular weight mucins limit *S. aureus* USA300 survival over time. USA300 growth (A) in SCFM (red), SCFM + eDNA (yellow), SCFM + LMW Mucin (blue), and SCFM + eDNA + mucin (purple) at 37°C for 4, 8, 12, and 24-hours. USA300 labeled with green fluorescent protein (GFP) fluorescence (B) was measured at 4, 8, 12, and 24-hours in a plate reader. Correlation plot (C) of USA300 CFUs and GFP fluorescence at 24 hours (R2=0.9938). Two-way ANOVA was performed on USA300 CFUs. (*P ≤ 0.05, **P < 0.01 ***P < 0.001). Data is represented by mean ± SEM.

Since media containing mucin is opaque, making it difficult to perform a standard growth curve based on optical density, we utilized our GFP-labeled USA300 strain to observe bacterial burden at these timepoints. The GFP fluorescence replicated results that measured viable CFUs, with in GFP fluorescence over time in non-mucin containing conditions (SCFM and SCFM with eDNA), while fluorescence in media containing LMW mucin remained stagnant (Figure 2B). We confirmed through a correlation plot utilizing CFUs from the 24 hour time point that levels of GFP fluorescence positively correlated with higher CFU counts (R^2^ = 0.9919; Figure 2C). To confirm the effects of LMW mucins on another strain of MRSA, we repeated these experiments using USA100, a hospital-acquired MRSA strain. We observed that USA100 also had a growth defect when in the presence of LMW mucin compared to HMW mucin (Figure S1A). Additionally, LMW mucin caused a significant decrease in CFUs of USA100 as early as 4 hours. At 24 hours, LMW mucin decreased GFP fluorescence in mucin containing conditions (Figure S1B-D). These results indicate that LWM mucin has a negative growth impact on both community and hospital-acquired *S. aureus* MRSA strains.

Although we saw eDNA could not recover *S. aureus* survival in the presence of LMW mucins, we wanted to confirm if the addition of eDNA changed other aspects of *S. aureus* biofilms, such as overall biomass or aggregate size. Microscopy of GFP-expressing USA300 showed no statistical difference in biomass of biofilms in grown in SCFM without polymers or SCFM with eDNA, and *S. aureus* formed confluent biofilms in both conditions (Figure 3A and B). As expected, LMW mucin significantly decreased biomass volume in media with and without eDNA (Figure 3A and B). Additionally, the average aggregate area was not found to be affected by the presence of both eDNA and LMW mucin compared to LMW mucin alone (Figure 3C). We repeated this assay using our GFP-expressing USA100 strain and saw similar results (Figure S2).

**Figure 3:**
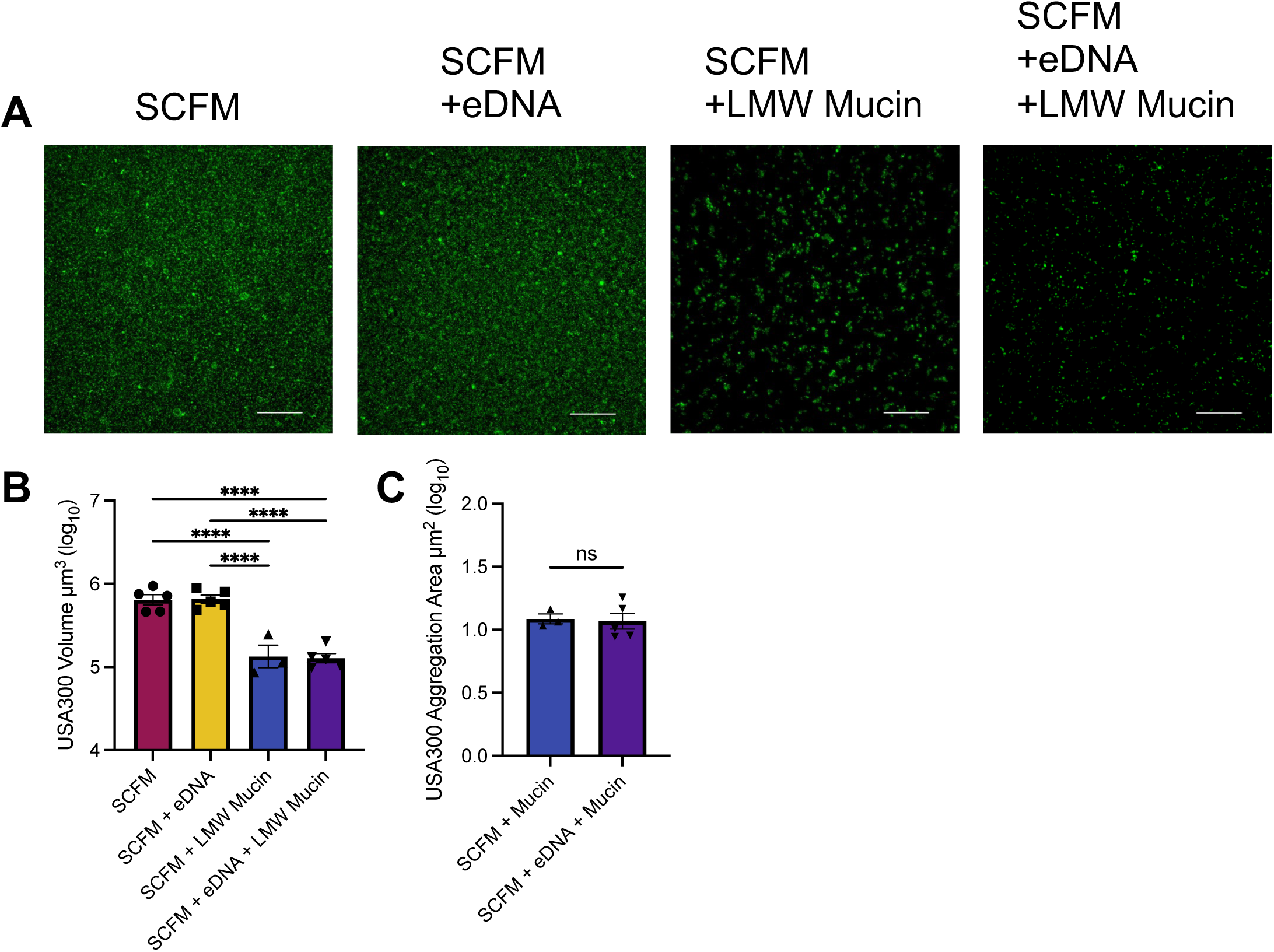
*S. aureus* USA300 biofilm biomass is reduced in the presence of low molecular weight mucin. Representative Z-stack images of USA300 (A) biofilm in SCFM with and without eDNA or LMW mucins after 24 hours. USA300 biomass (B) in each condition and average aggregate size in SCFM with LMW mucin with and without eDNA of 3-5 biological replicates and 5 images taken of each condition. One-way ANOVA was performed on volumetric measurement (\*\*\*\**P* ≤ 0.0001) and Welch’s t-test performed on aggregation area (ns = nonsignificant). Data is represented by mean ± SEM.

### Reducing LMW mucin concentration partially recovers *S. aureus* growth and increases biomass

Next, we wanted to determine if reducing the amount of LMW mucin would recover USA300 growth and biofilm biomass. We inoculated GFP USA300 into SCFM with LMW mucin at either 5 mg/mL or 2 mg/mL, representing mucin concentrations in CF or healthy airways, respectively (1), for 24 hours. Our images showed larger aggregates in 2 mg/mL LMW mucin compared to the higher 5 mg/mL condition representative of CF (Figure 4A). Through quantification of images, we confirmed that there was a significant increase of both biomass and aggregate size in 2 mg/mL LMW mucin compared to 5 mg/mL LMW mucin (Figure 4B and C). Next, we plated to measure CFUs and found that incubation in 2 mg/mL of LMW mucin resulted in a significant increase in USA300 survival compared to 5 mg/mL of LMW mucin (Figure 4D). However, there was still a significant decrease overall in USA300 survival compared to SCFM with no mucin. We repeated these experiments with USA100 and saw that USA100 also had a significant increase in biomass, aggregate size, and survival in 2 mg/mL LMW mucin compared to the 5 mg/mL (Figure S3). Together, these data indicate that reducing the amount of LMW mucin to concentrations found in healthy individuals improves *S. aureus* survival, biomass and aggregate size.

**Figure 4:**
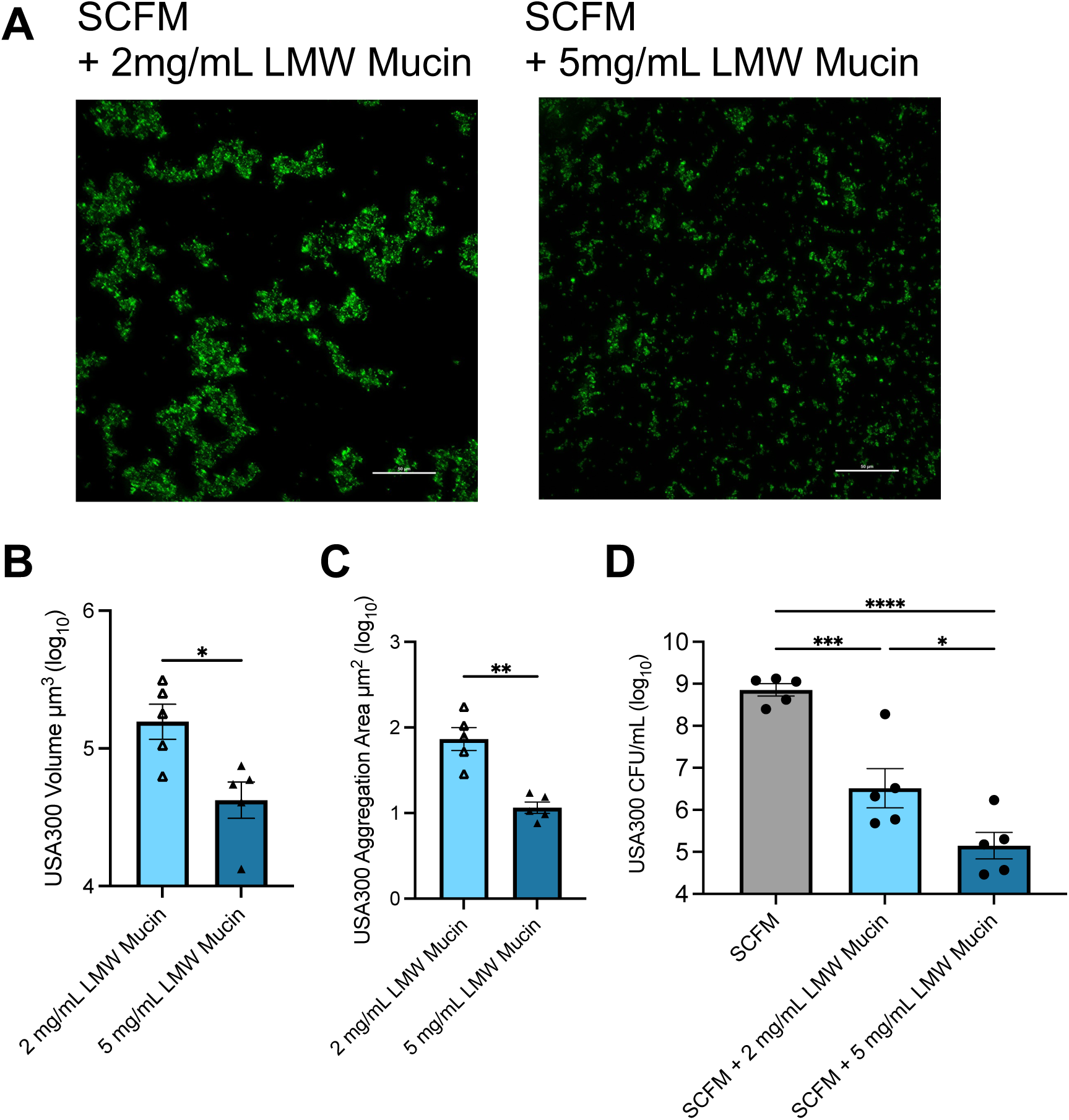
Reduced mucin concentration increases biomass and aggregate size and partially restores *S. aureus* USA300 survival. Representative z-Stack images of USA300 (A) in SCFM with either 2 mg/mL mucin (Left) or 5 mg/mL mucin (Right) after 24 hours at 40x. Scale bar represents 50 µm. Quantification of (C) volumetric and (D) aggregate size of 5 biological replications with 3-6 images taken of each well. CFUs of USA300 (D) after 24 hours of growth. One-way ANOVAs were performed on CFUs and Welch’s t-tests were performed on volume and aggregate area (*\*P* ≤ 0.05,*\*\*P* ≤ 0.01, \*\*\**P* ≤ 0.001, \*\*\*\**P* ≤ 0.0001). Data is represented by mean ± SEM.

### LMW mucin broadly impedes growth of common *S. aureus* laboratory strains and clinical isolates

Since we observed that LMW mucin results in a decrease of growth in two well-characterized MRSA strains, we next examined if this phenotype is broadly conserved. We evaluated additional *S. aureus* laboratory strains and *S. aureus* CF clinical isolates, including eight collected from the sinuses of individuals with CF and chronic rhinosinusitis (CRS) and one from the lung (Table S1). We utilized the fluorescence-based growth assay described above in Figure 2 to screen *S. aureus* strains transduced with the GFP-expression plasmid pCM29 over a 24-hour period. First, we determined the level of SCFM and mucin autofluorescence to ensure that all isolates were above the baseline level of autofluorescence. We measured all uninfected conditions and found that there was low autofluorescence that would not interfere with our bacterial reads (Figure S4). In SCFM without polymers, all laboratory strains and clinical isolates evaluated had an increase in fluorescence over time but varied in maximum fluorescence observed between strains (Figure 5A and E). The addition of eDNA to SCFM yielded similar results to SCFM without polymers (Figure 5B and F). As before, we saw that none of the laboratory strains or clinical isolates we tested had an increase in fluorescence over time in LMW mucin but did measure above our baseline levels of autofluorescence (Figure 5C and G). We continued to see this trend in SCFM with LMW mucin and eDNA for laboratory strains and clinical isolates (Figure 5D and H). Together, these data indicate that LMW has a broad negative impact across multiple laboratory and CF clinical isolates.

**Figure 5:**
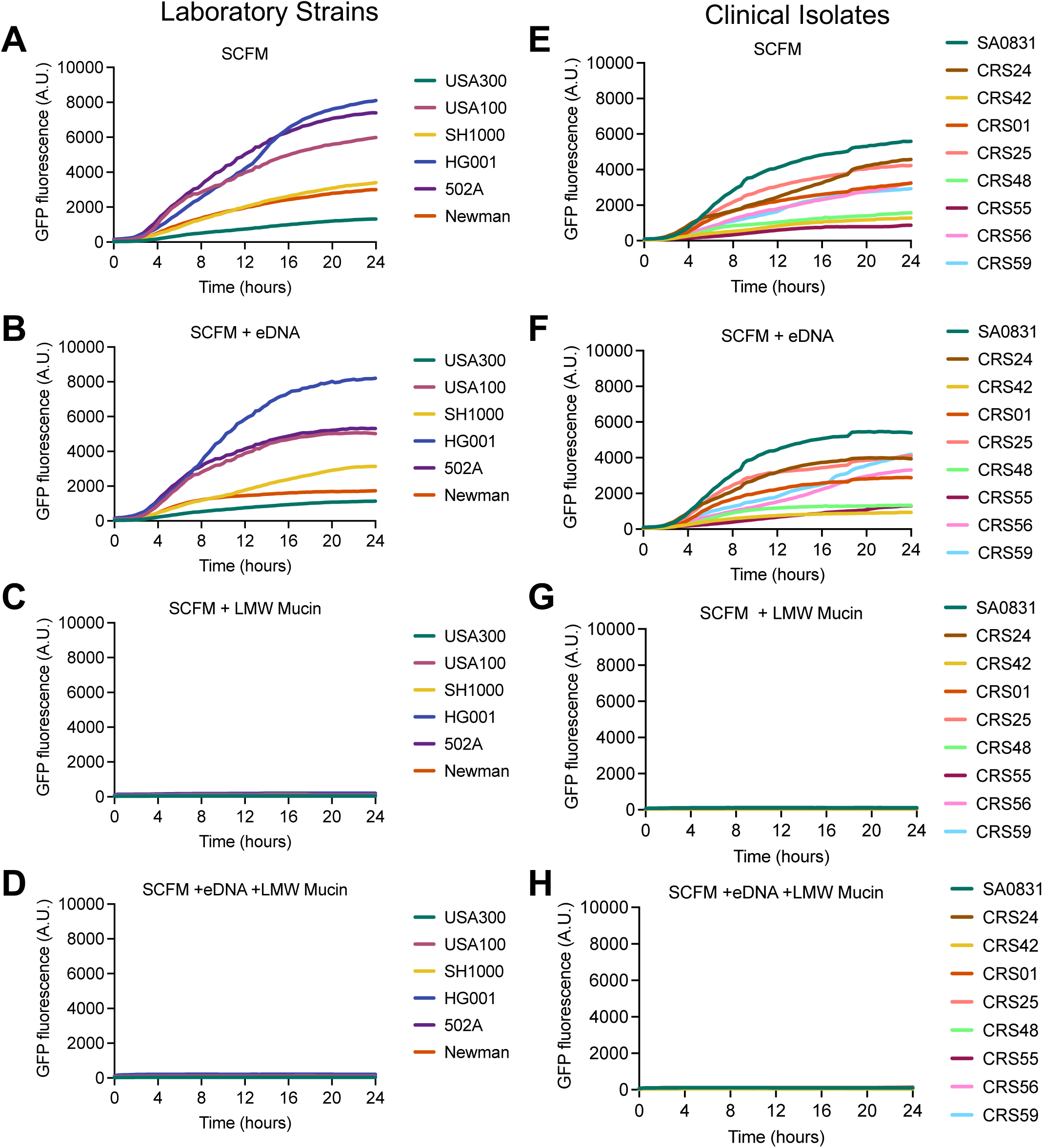
Mucin limits growth of *Staphylococcus aureus* laboratory strains and clinical isolates in SCFM. Laboratory isolates (A-D) and clinical isolates (E-H) of *Staphylococcus aureus* grown in SCFM conditions with GFP fluorescence reads every 20 minutes for a total of 24 hours. Lines represent a mean average of 5 biological replicates.

To determine if these strains can grow in the presence of LMW mucin in a different base growth medium, we dissolved 5 mg/mL of LMW mucin in tryptic soy broth (TSB). We first determined the baseline autofluorescence of uninfected TSB with and without LMW mucin (Figure S5A and S4). We then infected TSB or TSB with LMW mucin with the same panel of laboratory strains and clinical isolates. Like SCFM without polymers (Figure 5A and E), laboratory strains and clinical isolates exponentially grew in TSB (Figure S5B and D), however no strains tested were able to grow in TSB with LMW mucins (Figure S5C and E). This indicates that LMW mucins are mediating the observed decrease in growth and not the CF nutritional environment represented by SCFM.

### Investigating effects of LMW mucin on other CF pathogens and respiratory tract colonizing bacteria

Since we found that LMW mucin broadly impacts *S. aureus* strains, we next wanted to determine if other CF pathogens are similarly negatively impacted by LMW mucins. *Pseudomonas aeruginosa* is a common opportunistic pathogen in the CF respiratory tract, and mucoid variants of *P. aeruginosa* are associated with decreased lung function and quality of life in people with CF (30, 31). To test the effects of LMW and HMW mucins on *P. aeruginosa,* we utilized the non-mucoid and mucoid strains PAO1 and PAM57-15, respectively, inoculated into SCFM containing LMW or HMW mucin with or without eDNA. Both PAO1 and PAM57-15 grew approximately 2-logs above the inoculum in all conditions regardless of the presence of LMW mucin or HMW mucin (Figure 6A and B). PAO1 growth was significantly increased in SCFM containing HMW mucin alone compared to SCFM without polymers and in SCFM with LMW mucin and eDNA (Figure 6A). There was no statistical difference between conditions for PAM57-15 (Figure 6B). Next, we tested another gram-negative CF bacterial pathogen, nontypeable *Haemophilus influenzae* (NTHi). Compared to *P. aeruginosa,* NTHi maintained survival at the level of inoculum with no significant difference between LMW or HMW mucins with or without eDNA (Figure 6C).

**Figure 6:**
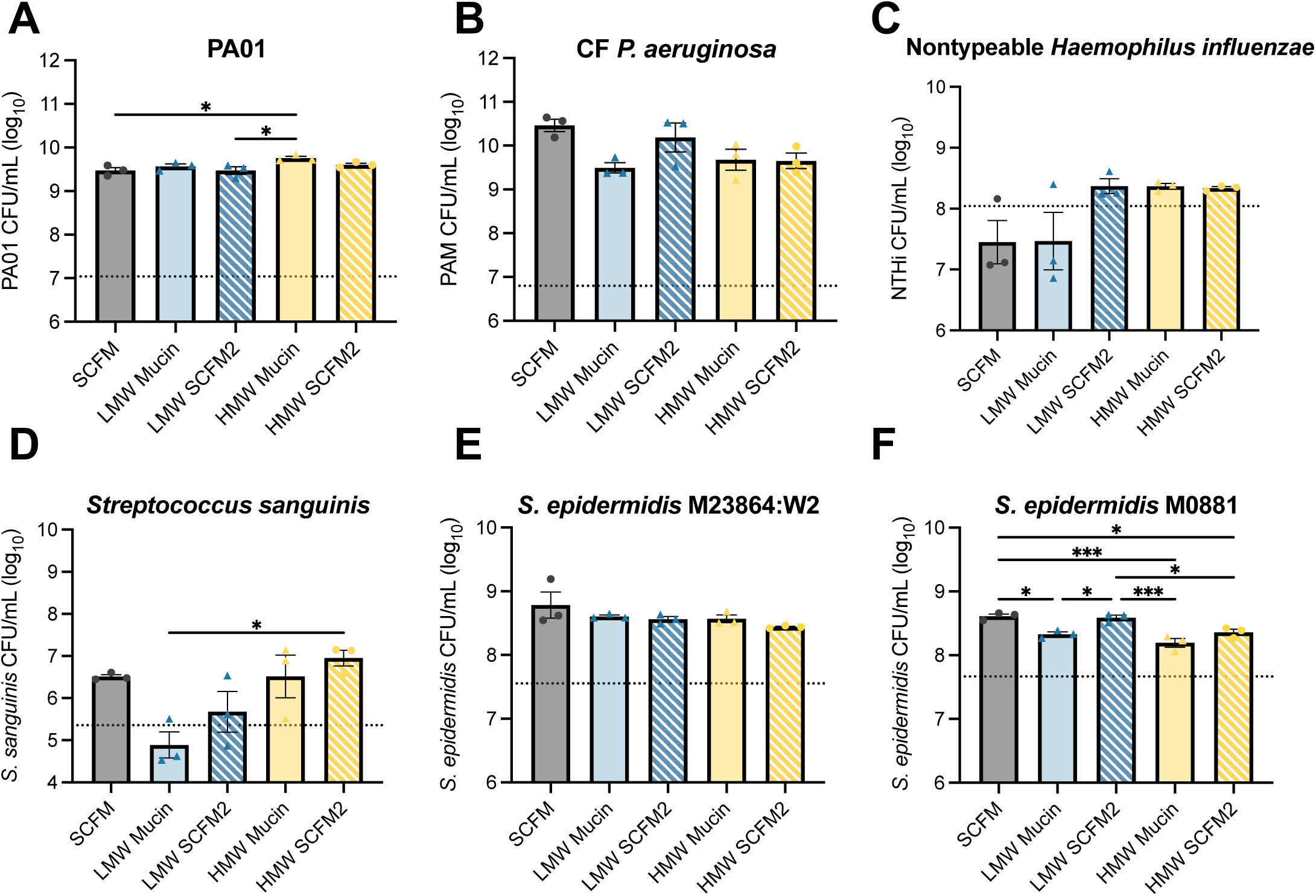
Other CF pathogens and *Staphylococcus epidermidis* are not negatively impacted by low molecular weight mucin. Growth of PAO1 (A), PAM57-15 (B), NTHi (C), *S. sanguinis* (D), *S. epidermidis* M23864:W2 (E), and *S. epidermidis* M0881 (F) in SCFM (grey), SCFM with LMW (blue) or HMW (yellow) with (striped) or without (solid) eDNA at 24 hours. One-way ANOVAs were performed on CFUs (*P ≤ 0.05,**P ≤ 0.01, ***P ≤ 0.001). Data is represented by mean ± SEM.

We next asked if other bacteria found in mucosal environments, such as the oral cavity, are impacted by mucins. *Streptococcus sanguinis* is an oral commensal species, and prior studies have shown that increased abundance of some *Streptococcus* species is associated with promoting stable lung function in pwCF (32, 33). While there was no significant difference in growth between SCFM without polymers and SCFM with LMW mucin, we observed a significant increase in *S. sanguinis* survival when grown in SCFM with HMW mucin and eDNA compared to SCFM with only LMW mucin (Figure 6D). To test if other *Staphylococcus* species are impacted by LMW or HMW mucins, we grew two strains of *Staphylococcus epidermidis* in the same conditions as above (Figure 6E and F). There were no significant differences between conditions with *S. epidermidis* M23864:W2 (Figure 6E). However, *S. epidermidis* M0881 had a significant decrease in growth in SCFM with LMW or HMW mucin compared to SCFM without polymers (Figure 6F). Additionally, there was a significant increase in CFUs when eDNA was added to the LMW mucin condition, but not the HMW mucin condition (Figure 6F). Despite these differences, both strains of *S. epidermidis* were able to grow at least a log above the inoculum, indicating that *S. epidermidis* is not experiencing the same negative effects of LMW mucin as *S. aureus* (Figure 1A and Figure 6E and F). Together, these results indicate that the growth inhibiting phenotype of LMW mucin may be specific to *S. aureus* and not to other species of bacteria that commonly inhabit the CF airways.

## DISCUSSION

Despite the introduction of HEMT to treat CF disease, bacterial infections, including those with *S. aureus*, are not eradicated despite the ability of HEMT to restore mucociliary clearance (21, 34, 35). In the work presented here, we showed that LMW mucins suppressed *S. aureus* growth and reduced survival when compared to HMW mucins (Figure 1 and Figure 2). In CF, increased levels of proteases released from innate immune cells and bacterial pathogens degrade intact airway mucins into smaller polymers (9, 10, 19, 25). These small mucin polymers can alter mucus biochemical properties by changing the elasticity and viscosity of the mucus layer, resulting decreased mucociliary clearance (25, 36). While there are some reported instances of bacterial degradation of mucus releasing secondary metabolites beneficial to *S. aureus* growth and persistence (19), our results indicate that without mucin-degrading bacteria present, LMW mucin has a inhibitory effect on *S. aureus*. Other studies that have inoculated *S. aureus* into SCFM2 have shown that genes related to iron acquisition are downregulated compared to human sputum samples, while genes related to virulence factors and metabolism are more closely comparable to human sputum samples (12, 37). However, these studies did not specify mucin polymer size used, and differential gene regulation may occur in the presence of LMW or HMW mucins. Further studies of transcriptional changes in low and high molecular weight mucin would be needed to understand how mucin polymers influence *S. aureus* gene expression.

*S. aureus* biofilm formation is largely governed by the accessory gene regulator (agr) system through the sensing of autoinducing peptides and production of extracellular dispersal molecules, such as nucleases, proteases and phenol soluble modulins (PSMs) (38). It has been found that eDNA released from cell lysis is important for *S. aureus* biofilm maturation and attachment (39–41). Our results also indicate that sputum polymers such as eDNA are not sufficient to rescue *S. aureus* from the negative effects of LMW mucins. In the experiments presented here, eDNA also did not provide a benefit when *S. aureus* was cultured in in HMW mucin. Interestingly, agr mutant *S. aureus* isolates are frequently identified from the respiratory tract, despite the known requirement of a functional agr system to produce secreted virulence factors and toxins associated with acute airway infection (42, 43). Further study of agr-regulated gene expression in the presence of LMW mucins, or potential defects in expression other biofilm-related genes, would be needed to determine if agr is dysregulated in the presence of LMW mucins. Alternatively, the negative impacts of LMW mucin on *S. aureus* may be dominant to any beneficial effects of eDNA on *S. aureus* biofilms.

Another group determined that *S. aureus* lacking the transcriptional regulator MgrA has a decreased binding affinity to both pig gastric mucin and BSM, but MgrA does not have the same affinity to other biopolymers (44). MgrA is a global regulator known to play a major role in biofilm formation, clumping, and repression of *S. aureus* cell wall-associated proteins (45, 46). MgrA is activated by the two-component system ArlRS, which regulates genes involved in agglutination and other virulence factors (45, 47). There is evidence that the *agr* system can control expression of *mgrA* through mRNA stabilization provided by the regulatory RNA, RNAIII (48, 49). *S. aureus* can also exhibit mucin-induced dispersal through production of PSMs, which are responsible for *S. aureus* biofilm dispersal as well as dispersal of other bacteria, including commensal Corynebacterium species as members of our group recently demonstrated (50, 51). It is possible that a combination of ArlRS, MgrA and agr-dependent signaling is needed to mediate survival in the presence of mucin through production of PSMs and expression of specific surface binding proteins.

Different muco-obstructive airway diseases can result in varying levels of mucin concentrations in the respiratory tract. The standard concentration of mucin in SCFM2 is 5 mg/mL, meant to replicate mucin concentrations observed in CF sputum from pwCF (1, 12, 24). Based on the literature, we wanted to replicate the average mucin concentration seen in the healthy population, which averages around 2mg/mL, for comparison to CF conditions (1, 25). As HEMT corrects ion transport across the epithelial layer in the CF airways, this has been found to result in restored mucociliary clearance, yet bacterial pathogens are often not eradicated. The continued persistence of bacteria such as *P. aeruginosa* in the HEMT-modified CF airway environment could result in continued degradation of mucins despite HEMT-mediated correction, leading to continued presence of small, pathogenic mucin polymer strands. Utilizing the LMW mucin at 2 mg/mL, we were able to show that as small mucin polymers concentrations decrease, *S. aureus* growth is partially restored, suggesting *S. aureus* could persist in this environment. Additionally, we show that S. aureus growth is not inhibited by HMW mucins, representing longer mucin strands that may be present in the HEMT corrected airway. Taken together, this provides some insight into why we see some bacteria, such as *S. aureus*, remain in the CF airways even after HEMT.

We observed that other CF pathogens are not negatively impacted by LMW or HMW mucin to the degree observed for *S. aureus* (Figure 6). Bacteria such as *P. aeruginosa* thrive in a mucus-rich environment and have developed ways to degrade mucin to benefit their growth and persistence (9, 52, 53). It was unsurprising to us that *P. aeruginosa* was able to grow in both LMW and HMW mucin (Figure 6A and B) due to previous reports of *P. aeruginosa* proteases, including LasB and an unidentified metalloprotease, that can degrade mucin strands (25, 54). NTHi does not have any known mechanisms for degradation of mucin polymer backbones, but it has a well-characterized sialidase that allows NTHi to utilize sialic acid residues from mucin side chains for biofilm maturation (55, 56). Interestingly, *S. epidermidis* was able to grow and persist in both LMW and HMW mucins, indicating that there are specific mechanisms that result in the inability of *S. aureus* to grow in the presence of LMW mucins.

Some limitations of this study include the inability to firmly control the size of the mucins in purchased BSM. Both BSM sources used contain a heterogeneous mixture of polymers of different lengths, with the HMW mucin having the largest range of polymer size. Additionally, other studies have noted that MUC5B monomer size averages about 671 nm in length, and polymers measure approximately 11 µm with overall molecular weight of ranging upwards of 40 mDa (57–59). Other groups have developed methods for collecting and purifying MUC5B from human saliva (60, 61). Utilizing native MUC5B mucin from human saliva in SCFM would give us a clearer understanding of how native mucins are impacting *S. aureus*. These interactions could be further characterized using a mucinase, such as StcE or bromelain, to cleave the mucin backbone into smaller polymer strands for further testing of *S. aureus* survival (62–64). Future studies would also benefit from examining differences in the mucin glycosylated side chains between the two available BSM sources used in this study and native MUC5B from human saliva to determine if a specific abundance of carbohydrate side chains negatively impacts *S. aureus* growth (8). Furthermore, our current model does not incorporate other antimicrobial factors that may be present in primary cell culture models, which will give important insights on how the epithelial layer may impact *S. aureus* survival (65).

## MATERIALS AND METHODS

### Bacterial Strains and Growth Conditions

*Staphylococcus aureus* species (Table S1) were grown on tryptic soy broth (TSB) with 1.5% agar (TSA; BD Bioscience) with or without chloramphenicol (10 µg/mL; Millipore Sigma) to maintain plasmids in GFP-expressing strains at 37°C. Overnight cultures were inoculated from a single colony in 5 mL of TSB shaking at 200 RPM at 37°C. For GFP strains, chloramphenicol (10 µg/mL final concentration) was added to maintain plasmids when necessary. Synthetic cystic fibrosis sputum media (SCFM) was made as described in Palmer *et al*. SCFM variations included bovine submaxillary mucins, high molecular weight (Thermo Scientific; J63859.ME) or low molecular weight (Millipore Sigma; M3895-1G), and/or UltraPure Salmon Sperm DNA Solution (Invitrogen) as indicated. For colony forming units (CFUs), overnight cultures of bacteria were diluted 1:1 in fresh TSB and spun at 4500 RPM to pellet bacteria. Bacteria were resuspended in of 1x phosphate-buffer saline (PBS; Fischer Scientific). Approximately 1 x 10^7^ bacteria were inoculated per well of a 96-well flat-bottom plate (Griener) containing 200 µL of SCFM with or without polymers. Plates were placed in an incubator, shaking at 37°C. CFUs were collected at 4, 8, 12, and 24 hours and plated on TSA plates. Fluorescence-based growth curves were performed in a Spark Automated Multimode Microplate Reader (Tecan).

### Transmission Electron Microscopy

SCFM with LMW or HMW mucin (5 mg/mL) were prepared and diluted to 50 µg/mL for TEM imaging. Samples were pipetted onto a carbon grid and allowed to incubate for one minute before washing with water and staining with 2% (w/v) uranyl acetate for one minute as previously reported (66). TEM images were acquired using a JEOL 1400 HC Flash microscope (JOEL USA).

### Fluorescence Microscopy and Quantification

Prior to inoculation with bacteria, 8-well glass-bottom chamber slides (Ibidi) were coated with 0.05% Poly-D-Lysine (MilliporeSigma) for 1 hour, washed three times with filtered MilliQ water and stored at 4°C until ready for use. 200 µL of SCFM was added per chamber, and approximately 1 x 10^7^ CFUs of GFP *S. aureus* was inoculated and allowed to grow statically for 24 hours at 37°C prior to imaging. Slides were imaged on a Ti Eclipse widefield fluorescence microscope (Nikon). Biomass and aggregate size were quantified using the Nikon NIS-Elements AR software package (Version 5.42.02 Build 1801). Biomass quantification was performed on image Z-stacks after thresholding and 3D deconvolving, and aggregate sizes were determined by the NIS-elements object count function on maximum intensity projection images. Data analysis of biomass and aggregate size was performed in Microsoft Excel (Version 16.96.1 [25042021]). Aggregates smaller than 1 µm were excluded from data analysis to account for noise. Images shown, biomass measurements and aggregate sizes are representative of 3-5 independent experiments with 3-6 individual fields of views measured for each sample for quantification.

### Statistical Analyses

Statistical analyses were performed with Graph Pad Prism version 10.4.2 (534) software (GraphPad by Dotmatrics). One-way analysis of variance (ANOVA) was performed on comparisons of CFUs at single time points, and two-way ANOVAs were performed on CFUs collected at multiple time points to determine significance between samples within a set time. Welch’s t-tests were performed on biomass and aggregate size. *P* values were considered significant if less than or equal to 0.05.

## ACKNOWLEDGEMENTS

Funding for this work was provided by Cystic Fibrosis Foundation awards DAVIS24P0 (Predoctoral training award to C.C.S.) and KIEDRO24P0 to M.R.K. Research reported in this publication (TEM) was supported by the UAB High Resolution Imaging Facility.

